# Evidence of latency reshapes our understanding of Ebola virus reservoir dynamics

**DOI:** 10.1101/2025.10.17.683141

**Authors:** John T. McCrone, Guy Baele, Ifeanyi F. Omah, Eddy Kinganda-Lusamaki, Joseph A. Brew, Luiz M. Carvalho, Nicola F. Müller, Gytis Dudas, Placide Mbala-Kingebeni, Marc A. Suchard, Andrew Rambaut

## Abstract

The reservoir of Ebola virus (EBOV) has remained a mystery since the first recorded zoonotic epidemic in 1976. While recent outbreaks have revealed much about the epidemiological dynamics that sustain human-to-human transmission, the mechanisms by which the virus persists between epidemics are unknown. Observations of extreme EBOV evolutionary rate heterogeneity in humans suggest latency is an important, yet untested, factor in persistence between epidemics. We here explicitly model latency with a novel phylogenetic approach to characterise the natural history of EBOV and, by extension, its reservoir. We find evidence that EBOV undergoes extended periods (i.e., possibly decades) of quiescence in the reservoir. Accounting for these dynamics significantly changes our understanding of EBOV’s evolutionary past and its geographic spread through Central Africa.

## Introduction

The first recorded outbreak of Ebola virus (EBOV) occurred in 1976 near Yambuku, in the north of what is now the Democratic Republic of the Congo (DRC) (*1*). The epidemic spread through the Yambuku Mission Hospital, resulting in 318 cases of Ebola virus disease (EVD) and 280 deaths. Because much of the transmission was nosocomial and the disease was unknown to people in the region, epidemiologists suspected the virus was introduced into the clinic by a traveller from Sudan, where a separate outbreak of haemorrhagic fever had recently been reported. However, it was noted at the time that EBOV may be endemic to the area and reside in an unknown reservoir. Anti-EBOV glycoprotein (GP) antibodies were found in five individuals who were not ill nor connected with the outbreak (*1*). Later, it was shown that the Sudan outbreak was caused by a separate filovirus – Ebola virus Sudan – further supporting a separate zoonotic origin of the outbreak in Yambuku (*2*). Since EBOV was first isolated in 1976, there have been at least 18 detected outbreaks sparked by independent zoonoses (see Table S1). It is now widely accepted that EBOV is endemic in the wildlife of Central Africa. However, the reservoir of EBOV remains uncharacterised.

EBOV is thought to be maintained among bat populations in Central Africa, based largely on observations that Marburg virus, a related filovirus, transmits among fruit bats (*3, 4*). Field work supports this hypothesis. Small fragments (∼ 265bp) of EBOV genomes as well as anti-Ebola GP antibodies were isolated from fruit bats sampled near human outbreaks along the border of the Republic of the Congo and Gabon in the early 2000s (*5*). Additionally, in 2019, EcoHealthAlliance sequenced a fragment of the virus from an insectivorous bat, the greater long-fingered bat (*Miniopterus inflatus*), in Liberia (*6*–*8*).

But the reservoir is likely more complex than one species or population. Partial sequences isolated from gorilla, chimpanzee, and duiker carcasses, and serology from non-human primates suggest multiple species act as intermediate hosts (*9*–*11*). The degree to which any particular species contributes to the long-term reservoir is unknown. Neither live virus nor whole genome has been isolated from a bat population, and concentrated sampling efforts near outbreaks have failed to find an active reservoir (*12*). Much of what is known about the EBOV reservoir has been derived from evolutionary analyses, which impute population dynamics from the ancestral relationships among zoonotic spill-overs (*11, 13, 14*). In the absence of any EBOV genome sequences from infected reservoir species, these approaches estimate the evolutionary dynamics of EBOV in non-human animals entirely from zoonotic infection of humans.

It has long been suggested that EBOV diversity dates back to the 1970s outbreaks of Yambuku (Yambuku/1976) and nearby Bonduni (Sud-Ubangi/1977) (*13*–*17*). The prevailing view relies heavily on a temporal rooting of the EBOV phylogeny estimated from isolates sampled between the Yambuku/1976 outbreak and the West African outbreak of 2013 (Makona/2013). Phylogeographic models based on this origin found EBOV exhibited a wave-like spread out of the northern DRC, and proposed an active expanding epidemic within the unobserved reservoir (*14*). The location of the Makona/2013 outbreak drew the geographic implications of this rooting into question, and led some to propose a new root position based on outgroup rooting with other *Ebola* species. This analysis found the West African outbreak as an outgroup to all other EBOV lineages (*18*). However, out-group rooting with divergent species is unreliable (*19*), and while the epidemic in West Africa was a geographic outlier (Figure S1A), its divergence from the 1976 outbreak in Yambuku is consistent with previous estimates of the virus’s evolutionary rate (Figure 1A) (*20*). Despite the unexpected location of the Makona/2013 outbreak, the temporally rooted model, with a root placement near the Yambuku/1976 outbreak, has persisted to this day.

**Figure 1:**
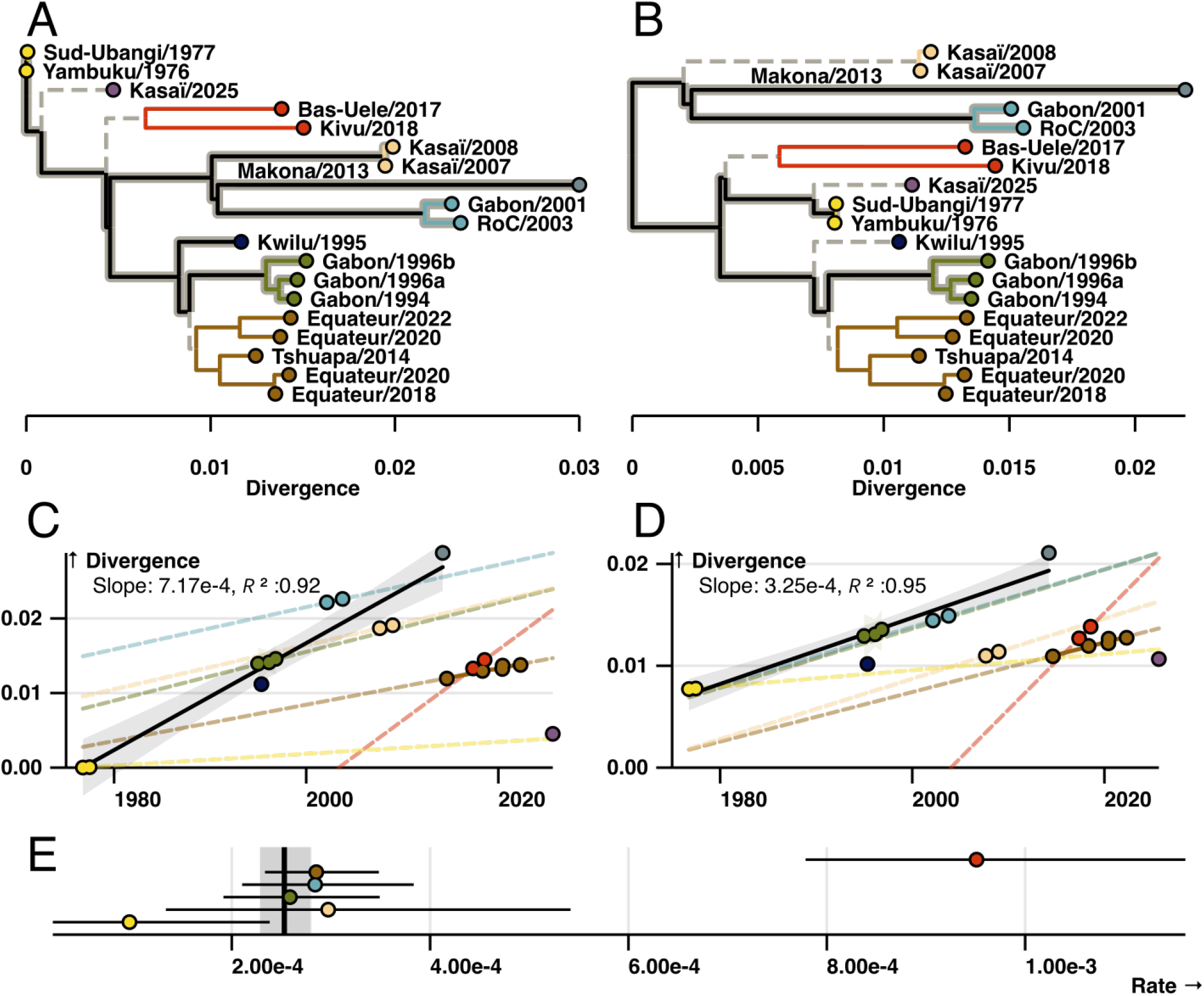
Evolutionary rate heterogeneity in the EBOV phylogeny. **(A & C)** The EBOV phylogeny rooted at the accepted 1970s outbreak position and corresponding root-to-tip divergence plot. **(B & D)** Similar to A and C but with the proposed rooting. In all figures tips are coloured by outbreak cluster (Figure S1A) and labelled by outbreak name (Table S2). Branches in A and B that are highlighted in grey connect tips used in the full regressions in C and D. Dashed, coloured regression lines are calculated for each cluster independently. **(E)** The maximum-likelihood evolutionary rate estimated for each cluster independently. Bars represent the 95% confidence interval (CI). The black vertical line represents the evolutionary rate estimated when all clusters share the same rate, with a grey box representing the 95% CI.

There have been eight sequenced zoonotic outbreaks of EBOV since 2013 (*21*–*25*) (Table S1). The most recent occurred in August, 2025 (Kasaï/2025) (*26, 27*). All have been less diverged than expected given the accepted model (Figure 1), leading some to hypothesize that EBOV may have entered a new reservoir population with slower replication dynamics (*21*). However, recent observations of EBOV rate heterogeneity in humans offer a different perspective. There are now multiple, well-documented cases in which EBOV has persisted in a recovered individual for extended periods of time (weeks to years), before sparking a secondary transmission chain (*28*–*31*). The maximum length of EBOV persistence in humans is unknown. The most extreme case to date lasted six years (*31*).

Genomic sequencing has revealed that fewer mutations accumulate during these long periods of persistence than expected given the EBOV evolutionary rate (*29, 31*). How infectious EBOV persists in a host for long periods without replicating (and by extension mutating) is unknown. The genomic epidemiology of these well-documented events implies it must.

We hypothesized that the decreased divergence of recent zoonotic events results from a latent process in the EBOV reservoir similar to that observed in persistent human infections, and that accounting for these dynamics would reconcile recent observations with the long-held model of EBOV’s evolutionary past. Here, we have developed a model of the virus’s evolutionary rate that accounts for periods of latency, and applied it to a dataset representing EBOV evolution in the reservoir. Our findings suggest EBOV latency is not unique to human infections, but is common throughout the natural history of EBOV. Furthermore, accounting for these dynamics draws the prevailing model of EBOV evolution into question and redefines our expectations for the reservoir behind zoonotic outbreaks.

## Results

We collected a representative set of Ebola virus genomes, taken from 19 plausibly independent zoonotic spill-over events (Table S2). The genomes included in this study were sequenced from human subjects, but because we restrict our sampling to one genome per outbreak and prioritize early samples, our phylogeny is largely shaped by evolutionary dynamics in the reservoir.

Restricting our dataset to genomes from the eleven recorded zoonotic outbreaks between 1976 and 2013, and rooting near the Yambuku/1976 outbreak recovers the accepted model of EBOV evolution and yields a root-to-tip regression with a high correlation coefficient and an R^2^ of 0.92 (Figure 1A & C). This rooting also produces an expected evolutionary rate of 7.17× 10^*−*4^ substitutions per site per year (s/s/y), not dissimilar to the 1.2 ×10^*−*3^ s/s/y (1.13,1.27 × 10^*−*3^) observed during human outbreaks (*32*) which is expected to be elevated due to incomplete purifying selection(*32, 33*).

However, as Lam *et al*. (*21*) and later Mbala-Kingebeni *et al*. (*34*) observed, this visually appealing linear relationship breaks down with the Tshuapa/2014 and subsequent outbreaks (Figure 1C). Attempts by us (*35*) and others (*34*) to account for the limited divergence of recent outbreaks have required topological constraints and/or informative priors on the molecular clock rate in addition to *a priori* assigned local clock models in order to recover a root near the original 1976 Yambuku outbreak. The necessity of these informative priors in addition to *a priori* assigned local clocks suggests the temporal signal in the first 11 outbreaks is not as strong as initially thought.

### Evidence for an alternative root position

It has been noted elsewhere (*34*) that the EBOV phylogeny is made up of several clades of outbreaks that cluster by time and location (denoted by colour in Figure 1 and S1). We observed that the accepted root-to-tip regression is driven by the long branches that connect these clusters (Figure 1A and C). However, there seems to be alternative regressions within each cluster (coloured clades in Figure 1), that differ consistently from this trend (Figure 1C).

We estimated an independent evolutionary rate for each cluster in Figure 1 using maximum likelihood and found remarkably similar rates across many clades (Figure 1E). Interestingly, this analyses suggested EBOV evolves slower within these clades (2.53 ×10^*−*4^) than expected given the accepted root-to-tip correlation. The clade representing the 2017 and 2018 Northern DRC outbreaks (Bas-Uele/2017 and Kivu/2018) may exhibit a higher rate of evolution. While this is consistent with previous results (*34*), we note that this clade is characterised by long external branches which corresponds to a high variance in the estimated rate.

We hypothesized that if the within-cluster evolutionary rate represents EBOV evolution in the reservoir then there should be a root position that allows for similar between-cluster dynamics. There is no rooting that does not require root-to-tip outliers with less divergence than their date would suggest. However, if we root the EBOV phylogeny as in Figure 1B, there is a subset of branches that pass through the root, connect several outbreak clusters, and imply a similar between-and within-cluster evolutionary rate.

We found statistical support for clock-like evolution with an identifiable rate in the subtree of eight outbreaks highlighted in Figure 1B when rooted at this proposed root (*36*). The proposed root position had the highest log-likelihood of all tested rootings (“Proposed” in Table 1), but a temporal signal was also observed when the tree was midpoint rooted with the Makona/2013 outbreak as an outgroup. Interestingly, we found no evidence for clock-like evolution in this dataset with the accepted root position near the Yambuku/1976 outbreak. These signals were maintained in a dataset comprised of the 11 outbreaks between 1976-2013 (“Classic temporal set”) as well as in the full dataset of 19 samples when local clocks were applied to branches subtending the remaining clades (dashed branches in Figure 1B; Table 1). Our findings suggest the accepted root position of EBOV is based on a misguided interpretation of root-to-tip correlations. However, clock-like evolution with periodic rate heterogeneity remains a plausible explanation for EBOV evolutionary dynamics.

**Table 1:** Temporal signal in subsets of EBOV outbreaks. Model - the clock model used (DR: different rates, i.e. no clock; SR: single rate, i.e. unidentifiable clock; SRDT: single rate dated tips, i.e. identifiable clock; localSR/SRDT - an SR/SRDT model with local clocks on dashed between-cluster branches Figure 1). For the 1970s rooting, branches that appear dashed in Figure 1A have local clocks. For all other roots dashed branches in Figure 1B are used. LL - Log likelihood, ΔLL - Log likelihood difference from the best model, df - degrees of freedom in a chi-squared test, p - p-value of a chi-squared test, Base rate - the inferred evolutionary rate (where applicable), Root - the root position used in the analysis (see Figure S1B)

| Data set | Model | LL | ΔLL | df | p | Base rate | Root |
| --- | --- | --- | --- | --- | --- | --- | --- |
| Temporal set<br>8 taxa | DR | -29893.4 | — | — | — | — | — |
| | SRDT | -29898.0 | -4.6 | 5 | 0.15 | $3.10 \times 10^{-4}$ | Proposed |
| | SRDT | -29900.3 | -6.9 | 5 | 0.02 | $1.96 \times 10^{-4}$ | Midpoint |
| | SRDT | -29906.6 | -13.2 | 5 | < 0.001 | $7.56 \times 10^{-4}$ | 1970s |
|  | SR | -29915.6 | -22.2 | 6 | < 0.001 | — | Midpoint |
|  | SR | -29931.2 | -37.9 | 6 | < 0.001 | — | Proposed |
|  | SR | -30036.9 | -143.5 | 6 | < 0.001 | — | 1970s |
| Classic temporal set<br>11 taxa + Local clocks | DR | -31462.5 | — | — | — | — | — |
| | localSRDT | -31466.6 | -4.1 | 6 | 0.32 | $3.05 \times 10^{-4}$ | Proposed |
| | localSRDT | -31469.8 | -7.4 | 6 | 0.02 | $2.02 \times 10^{-4}$ | Midpoint |
|  | localSR | -31487.9 | -25.5 | 7 | < 0.001 | — | Midpoint |
| | SRDT | -31503.0 | -40.6 | 8 | < 0.001 | $6.89 \times 10^{-4}$ | 1970s |
|  | localSR | -31504.0 | -41.6 | 7 | < 0.001 | — | Proposed |
|  | SR | -31634.6 | -172.2 | 9 | < 0.001 | — | 1970s |
| Full data set<br>19 taxa + Local clocks | DR | -34246.8 | — | — | — | — | — |
| | localSRDT | -34251.0 | -4.2 | 11 | 0.76 | $3.02 \times 10^{-4}$ | Proposed |
| | localSRDT | -34255.6 | -8.9 | 11 | 0.12 | $2.24 \times 10^{-4}$ | Midpoint |
|  | localSR | -34284.7 | -38.0 | 12 | < 0.001 | — | Midpoint |
|  | localSR | -34298.9 | -52.1 | 12 | < 0.001 | — | Proposed |
| | localSRDT | -34299.3 | -52.6 | 13 | < 0.001 | $6.02 \times 10^{-4}$ | 1970s |
|  | localSR | -34566.8 | -320.1 | 14 | < 0.001 | — | 1970s |

### A mechanistic rate model discovers novel evolutionary dynamics

We sought to determine if our proposed model for EBOV evolution is consistent with latency in the reservoir population. To this end, we developed a state-dependent evolutionary rate model in which lineages transition between latent and replicating states (see Materials and Methods). When in the replicating state, substitutions accumulate across branches according to an underlying evolutionary rate; however, the rate is zero whilst in the latent state. We constrain our model such that all nodes are in the replicating state. External nodes represent sampled, active infections, and internal nodes represent bifurcation events - both processes require replication. Because of this constraint, latency is less likely on branches that span short periods of time, such as those seen within outbreak clusters, and more likely on the longer branches that connect outbreaks.

We initially focused on the 18 outbreaks identified in Central Africa, based on the assumption that these may be connected by a contiguous reservoir population that excludes the geographically distant 2013 West African outbreak. The root proposed above had the highest posterior support (70.0%) of all root positions with the remaining support split between trees with the Gabon/2001-RoC/2003 clade or the Kasaï/2007-Kasaï/2008 cade as outgroups (Figures 2 & S2). This analysis resulted in a mean root age of 1929 (95% HPD: [1887, 1957]) which corresponded to a mean replicating evolutionary rate of 2.34 ×10^*−*4^ s/s/y (95% HPD: [1.41× 10^*−*4^, 3.19 × 10^*−*4^]) consistent with the exploration above. We find evidence for wide-spread latency with median of 6 branches exhibiting at least one period of latency (95% HPD: [5, 9]) (Figure 2A). The average duration of latency on a branch with at least one period of latency was 21.7 years (median 20.6 years) (Figure 2). The five branches observed to be outliers in the root-to-tip plots are the only branches with a greater than 50% posterior probability of at least one latent period (Figure 2A).

**Figure 2:**
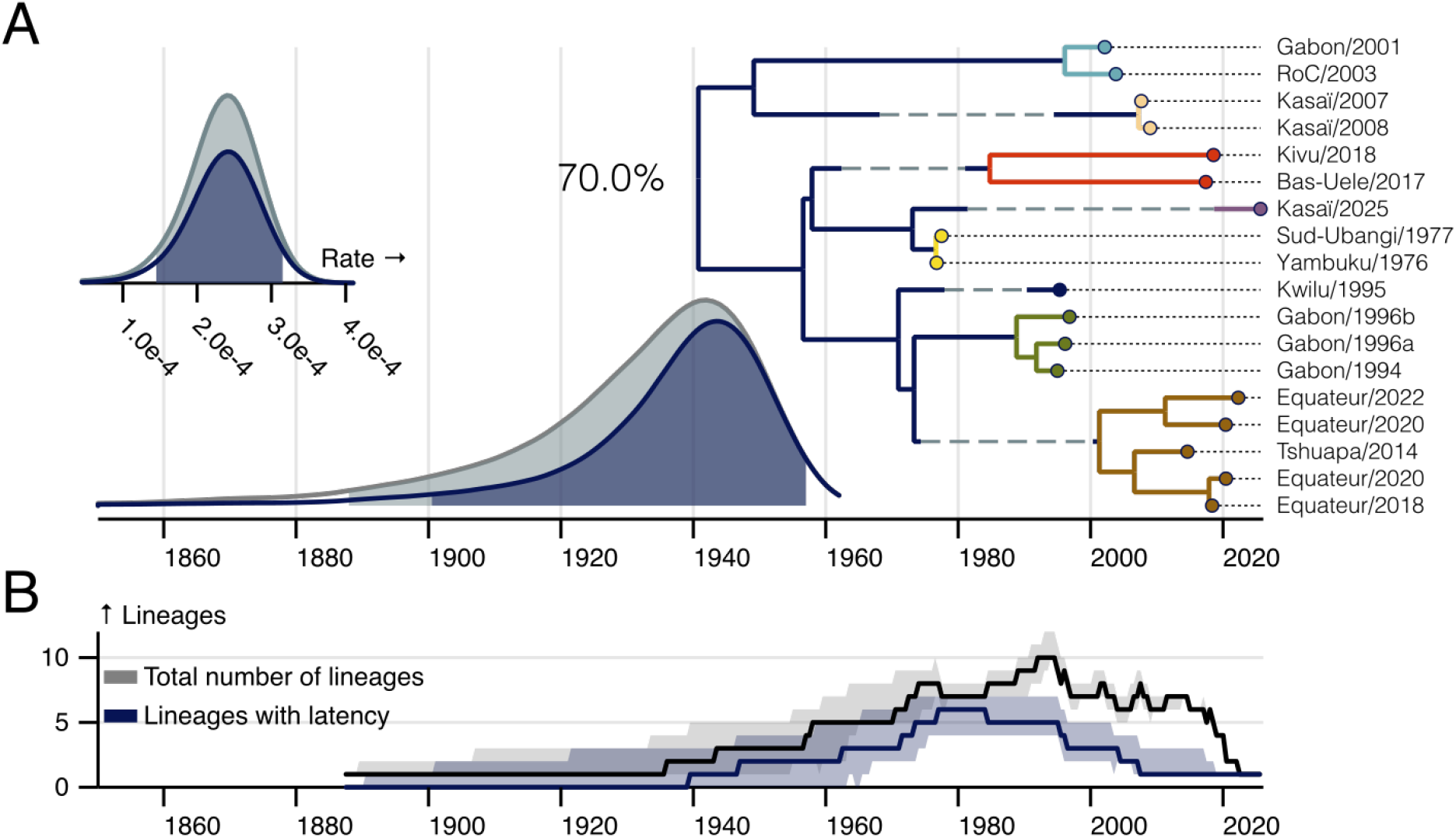
The evolutionary history of EBOV in Central Africa under the latent model. **(A)** A maximum clade credibility (MCC) of tree of 18 EBOV outbreaks in Central Africa. The tree summarises all trees with the displayed rooting. The posterior support for this rooting is displayed next to the root. Clades are coloured as in Figure 1 with tips labelled by outbreak. The root age distribution of the entire analysis is shown in grey with the contribution of the displayed root highlighted in blue. For branches with greater than 50% posterior support for latency, the posterior probability of latency is labelled above the branch. The mean duration of latency (conditioned at least one period of latency) is shown as a dashed line centered between the parent and child node. The marginal posterior root age distribution is shown in grey with contribution of the displayed root highlighted in blue. Inset: The posterior distribution of the evolutionary rate during replication, shaded by total analysis (grey) and conditioning on the displayed root (blue). In all posterior distribution plots, the shaded area represents the 95% HPD. **(B)** The median number of lineages that show any latency across the entire posterior is plotted a blue solid line, with the 95% HPD shown as a shaded area. The median total number of lineages through time and corresponding 95% HPD is plotted in grey.

Our analysis suggests all latent lineages were seeded prior to 1980. These lineages persisted until the mid 1990s, when they sparked a second wave of diversification (Figure 2B). Interestingly all zoonotic outbreaks observed after the year 2013 derive from once latent lineages (Figure 2A).

Despite being a geographic outlier, the 2013 West African outbreak is not a temporal outlier (Figure 1D). Is a branch of this length consistent with our estimates of latency? If we assume this outbreak diverges from the rest of EBOV near the root estimated above, its expected branch length would be 85 years long (95% HPD [57, 127]) which corresponds to a 4.96% probability (0.62%, 19.22%) of no latency.

Our findings were robust to the inclusion of this possible outlier, with the exception of increased uncertainty in the EBOV root placement. The mean root age of the extended dataset was 1918 (95% HPD [1872, 1954]) which corresponded to a similar replicating evolutionary rate of 2.34 ×10^*−*4^ substitutions per site per year (95% HPD: [1.50 ×10^*−*4^, 3.13 × 10^*−*4^]). Inclusion of the Makona/2013 outbreak resulted in a median of 7 branches with latency (95% HPD: [5, 10]), with similar lineage through time dynamics (Figure 3C).

**Figure 3:**
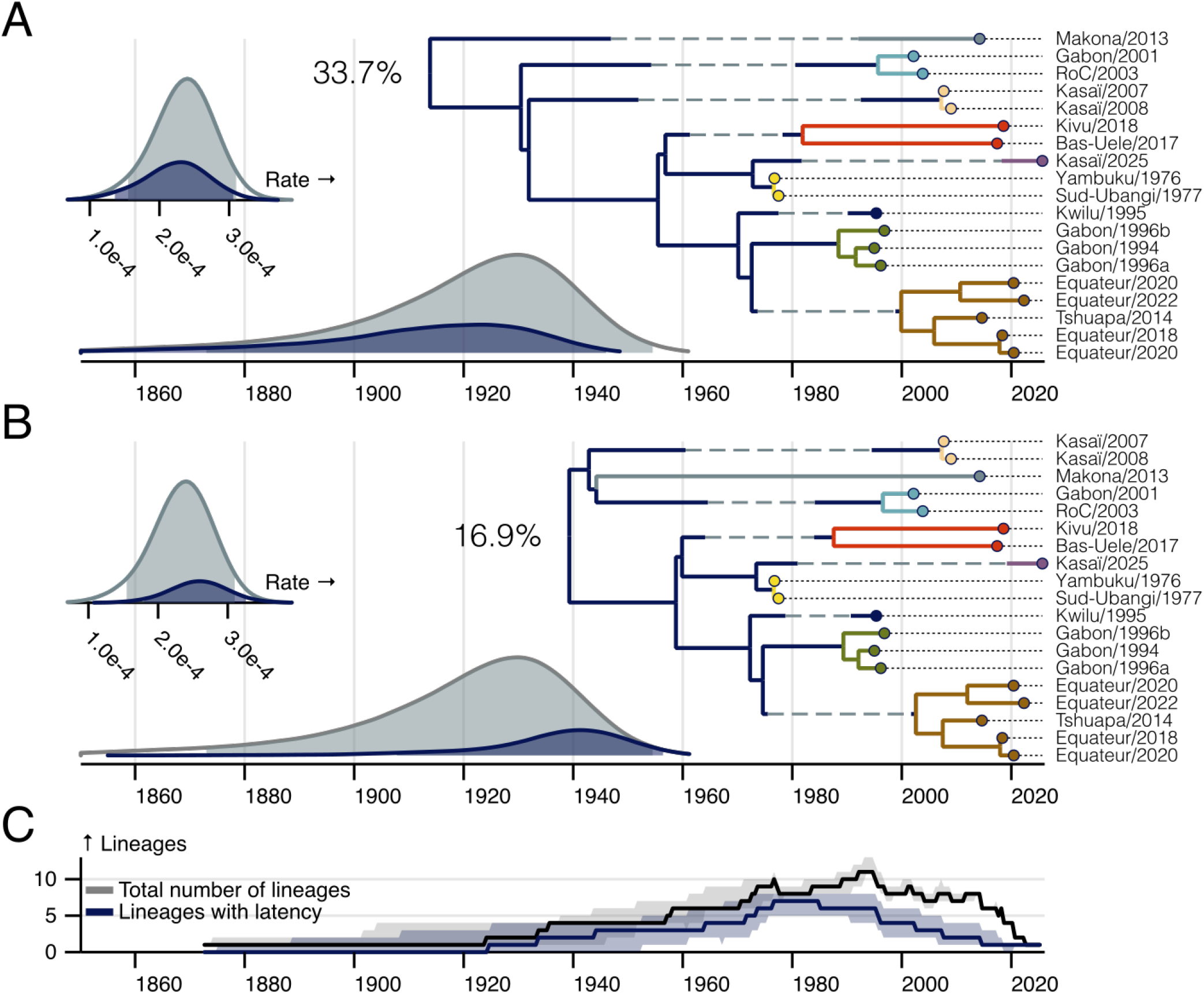
The evolutionary history of all EBOV outbreaks under the latent model partitioned by root position. **(A & B)** MCC trees of the most common root placement (A) and the proposed root (B) (posterior probability noted near root). Clades are coloured as in Figure 1 with tips labelled by outbreak. The marginal posterior root age distribution is shown in grey with each root’s conditional contribution is highlighted in blue. For branches with greater than 50% posterior support for latency, the posterior probability of latency is labelled above the branch. The mean duration of latency (conditioned on there being at least one period of latency) is shown as a dashed line centred between the parent and child node. Insets: The posterior distribution of the evolutionary rate during replication, again shaded by total analysis (grey) and conditioning on specific roots (blue). In all posterior distribution plots the shaded area represents the 95% HPD. **(C)** The median number of lineages that show any latency across the entire posterior is plotted a blue solid line, with the 95% HPD shown as a shaded area. The median total number of lineages through time and corresponding 95% HPD is plotted in grey.

The underlying evolutionary rate is consistent with previous estimates, but we find less support for the proposed root position (16.9% posterior support, Figure 3B). Instead, the model favours a root with the Makona/2013 outbreak as an outgroup (33.7%) (Figure 3A). The remaining posterior support was again split between trees with the Gabon/2001-RoC/2003 clade or the Kasaï/2007-Kasaï/2008 clade as outgroups (Figure S3).

### The geographic spread of EBOV in Central Africa

Previously, it was suggested that EBOV emerged from a genetic bottleneck near the Yambuku/1976 outbreak and then rapidly spread west through a wave-like process with a mean wave-front velocity of approximately 50 kilometers per year (*14*). This estimated dispersal is a direct consequence of the root placement, and is inconsistent with our updated model.

We employed a continuous phylogeographic diffusion model with a single diffusion rate to characterize the geographic dispersal of EBOV alongside the latency model of EBOV evolution. It has recently been show that more flexible phylogeographic approaches, which model diffusion as a branch-specific process are prone to misleading conclusions and overconfidence when applied to dynamic populations (*37*). Although our model is likely over-simplified, it provides an interpretable starting point to explore the geographic implications of latency in the EBOV reservoir.

We found the EBOV root was located in western DRC near the 2014-2022 outbreaks in the Equateur province. This location is not unexpected given the nature of the phylogeographic diffusion modeled, and the topology of the tree, which is likely to place the root between geographically dispersed outbreak clusters (Figure 4A and 4B). Contrary to previous models, which found EBOV spread directionally from its root location, our model suggests EBOV dispersal was initially limited. The location of most ancestral nodes matched that of the root (Figure 4B).

**Figure 4:**
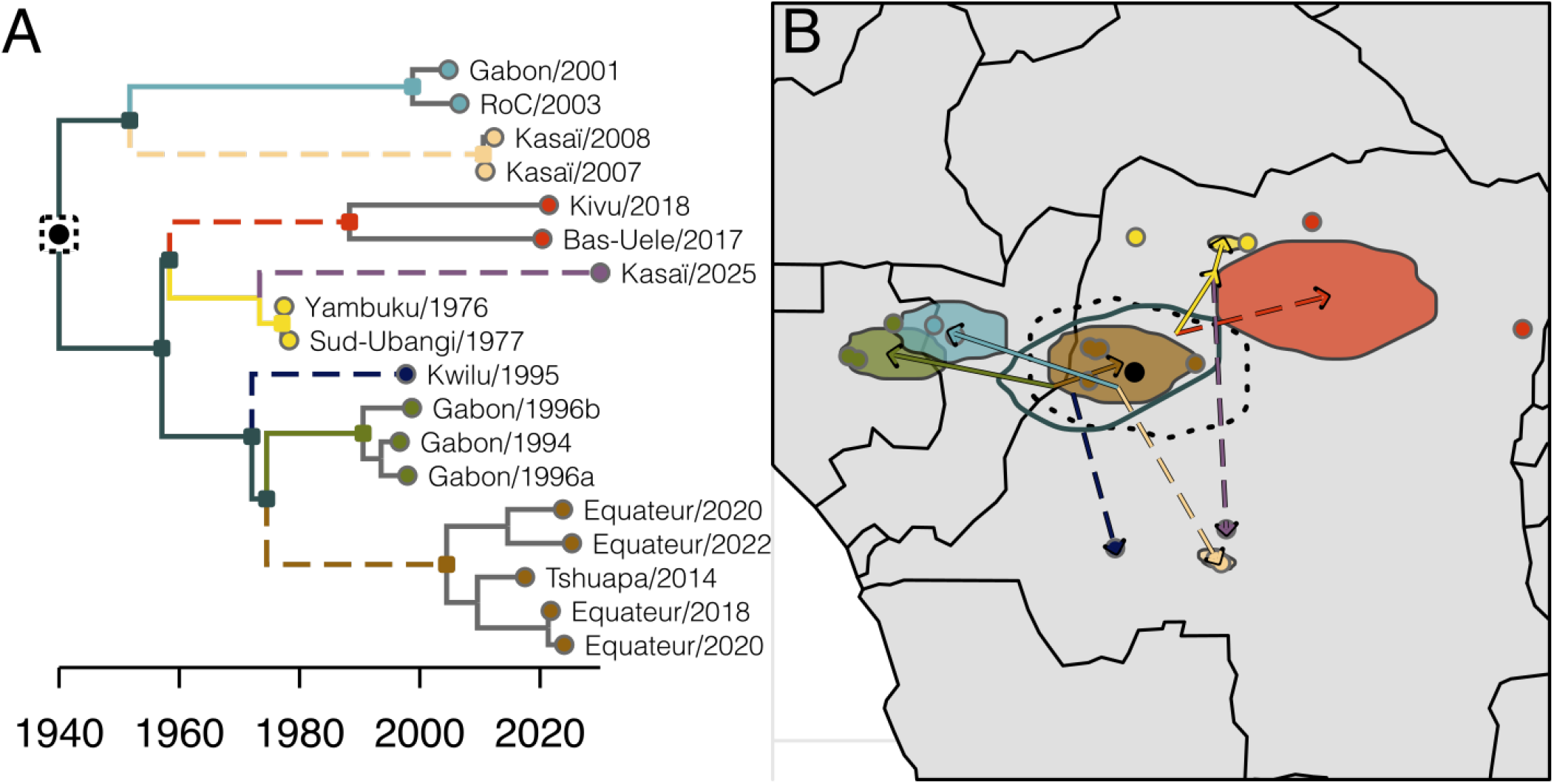
The geographic implication of latency in the EBOV reservoir. **(A)** The MCC tree of the dataset when the West African outbreak is excluded. Branches with more than 50% posterior probability of at least one latent period are dashed, and those present in the map in B are coloured according to the clade they subtend. Tips are labelled by outbreak and coloured as previously. The root is denoted with a black circle surrounded by a dashed box, while ancestral nodes shown in B are highlighted with dark squares. Clade roots are highlight with a coloured square. **(B)** A map depicting the geographic spread of EBOV. The average root position is represented by a black circle. The 50% HPD of the root position is enclosed by a dashed contour, while the 50% HPD locations of the ancestral nodes (squares in A) is represented by the solid contour. The arrows represent the mean source and destination of the similarly coloured branches in A. Outbreaks are shown as coloured circles, and the 50% HPD of the root of each outbreak clade is denoted as a shaded contour.

The location of the outbreak clusters and tree topology requires migrations from this ancestral source along three different axes (one northwesterly, one southerly, one northeasterly). Interestingly, multiple independent lineages – some latent – follow each of these corridors. Similar geo-graphic histories are not expected in independent clades of the tree in a purely diffusive process where location and genetic distance are highly correlated. These observations are robust to the inclusion of the West African outbreak which shifts the root location to the west and provides an addition migration along the northwestern axis (Figure S4). Together, these analyses suggest a ge-ographic model in which EBOV circulation was constrained near the recent outbreaks in Equateur, DRC, followed by repeated disseminations along several migratory routes.

### The plausibility of future latent-derived outbreaks

Our results suggest a significant number of EBOV outbreaks have derived from once latent lineages, and that latency may be an evolutionary mechanism by which EBOV persists in the reservoir. To determine the contribution of latency to future EBOV outbreaks, we plotted the cumulative distribution of there being at least one latent period as a function of branch length using the parameters estimated by our model (Figure 5A). We excluded the West African outbreak in these parameter estimates and relied on the analysis presented in Figure 2 on the basis that the West African outbreak seems to have originated from a separate process than that acting in Central Africa. Our results suggest a branch length of 35 years has a 50% chance of exhibiting latency, and the expected proportion of time spent latent for such a branch is 0.4 or roughly 14 years. As highlighted in Figure 5B, we have little precision in our prediction of the sum duration of latency, provided there is latency on a branch.

**Figure 5:**
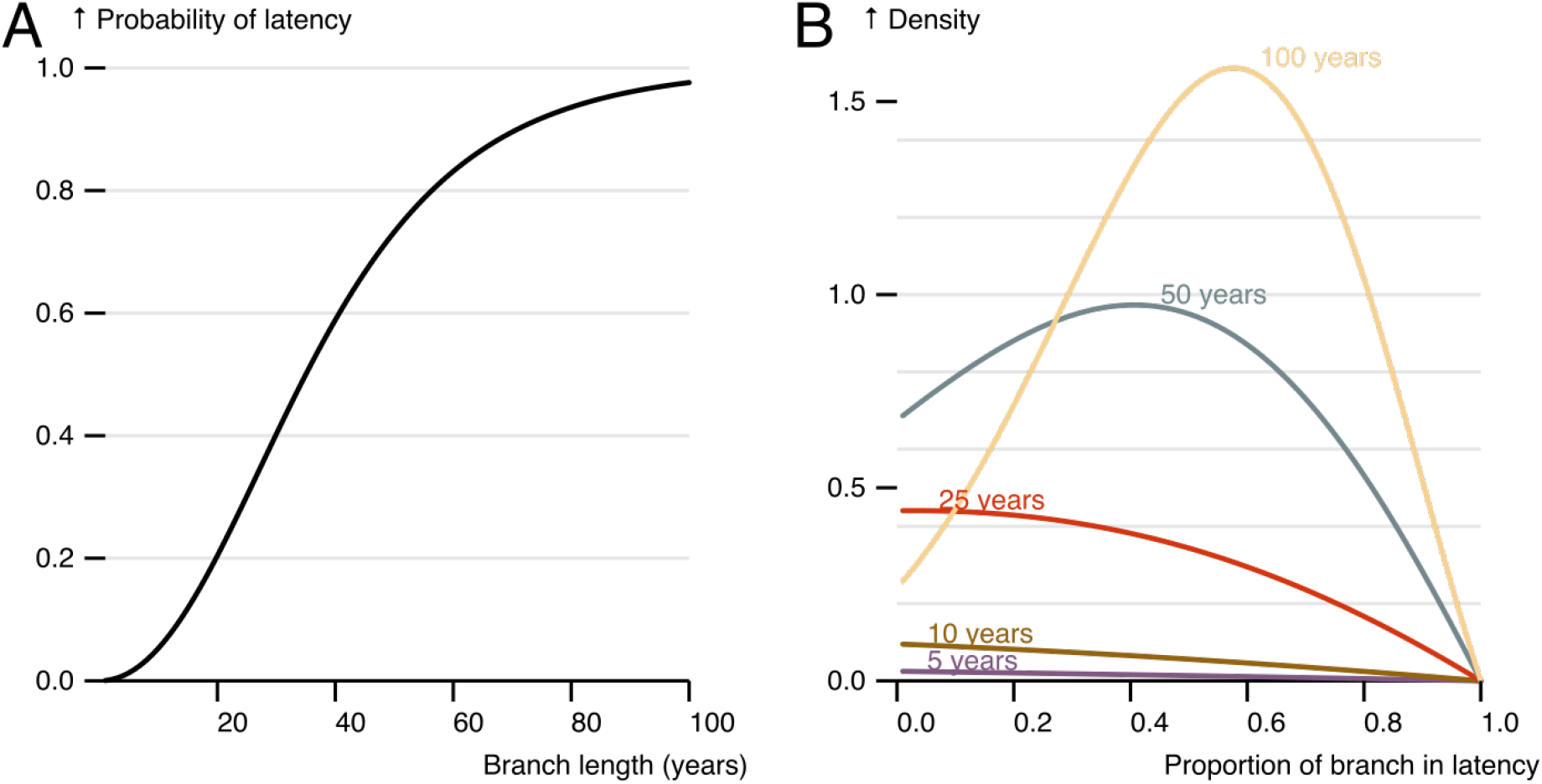
The expected contribution of latency to future outbreaks based on model estimates. **(A)** The probability of some latency as a function of branch length. **(B)** The distribution of the proportion of time spent in latency provided there is some latency on branches of different length (noted text near each line).

## Discussion

An accurate model of EBOV evolution in the reservoir has important consequences for how we classify outbreaks and evaluate the risk of future epidemics. Decreased rates of evolution are thought to be a marker of human-derived outbreaks (*38*). In the initial phylogenetic analysis of the Equateur/2020 outbreak, one genome was found to cluster near the base of the Equateur/2018 outbreak (*23*). The sample was thought to represent secondary transmission from a persistent human case. However, there was no epidemiological evidence for this noted at the time, and this sequence’s divergence is consistent with our modified estimate of EBOV’s evolutionary rate (note the two Equateur/2020 isolates in Figure 1). Given our updated model, this sample likely represents an independent spillover event from a large, diverse reservoir active in Equateur province, DRC, between 2018-2022, and not a persistent human case.

Given our finding that latency is wide-spread in EBOV evolution, it may be tempting to conclude that many EBOV outbreaks have resulted from persistently infected survivors of previous, unobserved outbreaks. Perhaps humans make up a significant proportion of the reservoir. We believe this to be implausible. Persistence in humans – although observed – is exceedingly rare, and only associated with the largest epidemics. Such an EBOV reservoir would require large, unobserved human epidemics. Additionally, our analysis identifies latency on several internal branches, precluding the possibility that these branches represent persistent human infections.

Our model suggests EBOV diversity predates the first observed outbreaks in the 1970s, and does not require EBOV to have emerged from a genetic bottleneck just prior to the first recorded human outbreak. Although seemingly supported by the root-to-tip model at the time, this bottleneck and subsequent rapid expansion across the continent defied ready explanation from a disease ecology point of view. We propose, instead, that EBOV spread across the region from an ancestral source outbreak located near the recent zoonoses in Equater, DRC. Future geographic reconstructions should explore whether this spread was gradual, or the result of a sudden dissemination perhaps due to changing land use practices in the 1960s and 70s.

Finally, we hypothesize that the phenomenon of EBOV persistence and latency in the reservoir is a mechanism by which the virus can maintain itself long term in a highly-structured host population - i.e., high-density but geographically dispersed bat roosts. Several of the migratory branches proposed by our phylogeographic model exhibit latency. Younger bats dispersing to other colonies for mating, whilst persistently infected, may be the route of transmission between roosts. Furthermore, persistent infections within a roost may spark a new outbreak years later after sufficient immunologically-naïve individuals have been born. This then establishes a new cohort of persistently infected individuals. While similar models of persistence have been proposed (*39, 40*), such hypotheses have remained untested. Here we have shown persistence combined with periods of latency is consistent with the phylogenetic patterns we observe between human outbreaks of EBOV.

The reservoir composition and dynamics of Ebola virus have evaded characterisation since the virus was first isolated in the late 1970s. Our work here has used phylogenetic analyses to explore the reservoir dynamics from its periodic spillover into humans. Filling in the details, the mechanisms behind latency, and its implications for host dynamics are crucial to understanding when and where EBOV will emerge next.

## Materials and Methods

### Data

To explore EBOV evolutionary rate variation in non-human hosts, we assembled a data set of genomes that span the known history of the virus. Most available EBOV genomes have been sampled from human cases. We have included one genome per outbreak, preferring those with precise dates of sampling. A list of sequences used is provided in Table S2, including, where applicable, their GenBank and Pathoplexus accession numbers. A SeqSet of all genomes use in this study can be found on Pathoplexus (*41*).

### Phylogenetic inference

Sequences were aligned using MAFFT (v7.526) (*42*) and refined by hand. Five prime and three prime untranslated regions were ignored in all phylogenetic analyses.

In each analysis, we used a separate HKY substitution model (*43*) for each of three partitions of the data: coding position one, coding position two, and a concatenation of coding position three and the intergenic regions. This was deemed to be the best partition scheme by IQ-TREE partition finder when compared to a partitioning scheme with separate partitions for each of the tree codon positions and intergenic regions. Maximum-likelihood phylogenetic trees were estimated using IQ-TREE v3.0.1(*44*). The likelihood ratio tests in the TipDate analysis (*36*) were performed using PAML (commit:ac9b97c8c35) (*45, 46*).

For the root-to-tip correlations in Figure 1, which exclude several clades, we first rooted the tree on the specified branch and then determined the location of the root on that branch based on most likely location from the TipDate analysis above. These calculations gave very similar results to those obtained from TempEst when run on trees containing just the tips included in the regressions (highlighted branches in Figure 1A and 1B) (*47*).

Clade-level, maximum-likelihood evolutionary rate estimates were calculated using the strict-clock model proposed by Didelot *et al*. (*48*). Each clade was identified in and pruned out of the full maximum-likelihood phylogenetic tree estimated by IQ-TREE so that even clades with only 2 tips were rooted. We then estimated the maximum-likelihood evolutionary rate and time-scaled the branch lengths using a gamma distribution with mean equal to the variance as described in (*48*). Optimization was achieved using the optimization library fmin (https://www.npmjs.com/package/fmin).

All Bayesian phylogenetic analyses were performed in BEAST X (*49*) using the BEAGLE high-performance computational library (*50*) to improve computational performance. For each analysis two chains were run for 500 million states and sampled every 100 000 states with the first 25 000 states removed as burn-in. We employed a constant population coalescent tree prior with a log-normal prior (logarithm of location *µ* = 4, logarithm of scale *σ* = 1) (95% interval [8, 388]) on the population size. Preliminary analyses with an exponentially growing population did not exhibit a growth rate different from zero. Phylogeographic analyses employed a time-homogeneous Brownian diffusive model of geographic spread (*51*).

We placed a conditional reference prior on the underlying evolutionary rate (*52*), with a slight modification. Typically, this prior is approximated as a gamma distribution with shape 0.5 and rate equal to the length of the tree, which results in an expectation of 0.5 mutations at each site over the entire tree. Because our tree contains periods during which no mutations accumulate, we updated the prior so that the rate was equal to the length of the replicating part of the tree which arrives at the same expectation.

We used a similar reference prior on the latent transition rate (see below for details). However, because latency is a branch-specific event that ‘resets’ at each node and because we expect there to be some small amount of latency over the duration of the tree, we use the root height as the rate parameter of the prior as opposed to the length of the tree. Sensitivity analyses with half the root height and twice the root height yielded similar results (Figures S6 & S7).

We place a uniform prior on the relative rate bias between entering and leaving the latent state. Sampling from these prior distributions and the samples dates of the full 19-taxa dataset results in a mean of 0.06 branches enjoying at least one period of latency with a 95% prior probability of no branches exhibiting latency. We assessed convergence to the posterior distribution and proper mixing of all relevant parameters in Tracer 1.7 (*53*).

All BEAST X XML files used in this analysis can be found at https://github.com/ebov/latency-reservoir.

### Latent-state branch rate model

We model the evolutionary rate of EBOV according to a state-dependent rate model (*54*) where branches alternate between two possible states, replicating and latent, according to a continuous-time Markov chain (CTMC). Each branch transitions independently between states according to an infinitesimal rate matrix Q conditioned on starting and ending in the replicating state. This condition is necessitated by the fact that each node represents either a sampled active infection (external nodes) or an unsampled branching event (internal nodes). While in the replicating state, each branch undergoes evolution according to some underlying molecular clock model (e.g. strict clock, uncorrelated relaxed clock, etc.); the evolutionary rate is set to 0.0 when in the latent state. The overall evolutionary rate along a branch is then determined by its underlying rate (*µ*) and the amount of time spent in the replicating (t_r_) and latent (t_l_) states, where t = t_l_ + t_r_ is the total length of the branch in time.

Several approaches exist for estimating the amount of time a branch spends in each state. Cappello *et al*. explicitly sample the state history of each lineage as it transitions between replicating and non-replicating states in a seedbank model (*55*). While accurate, explicitly sampling complete transition histories is computationally expensive. Similarly, Lewinsohn *et al*. have developed an alternative state-dependent evolutionary rate model that marginalizes over the state history and approximates the evolutionary rate of each branch by using the expected time spent in each state (*54*). Such an approximation seems appropriate in cases where both states replicate at different rates. However, in our setting – where branches may not experience any latency – the mean time spent latent is a poor representation of the possible dynamics.

We avoid sampling transition histories and instead estimate the proportion of time each branch spends in latency, integrated over all possible histories. Our approach is akin to recent approximations to the structured coalescent which also marginalize over histories (*56*–*59*). However, the model used here is simplified in that each branch’s state history is independent of all others, which allows for an exact solution.

We are interested in estimating the phylogenetic posterior distribution described by

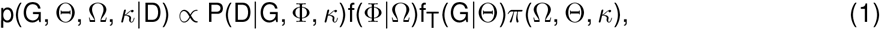

where G is our phylogenetic tree with branch lengths in years, Θ represents the parameters governing the coalescent tree prior, Ω the parameters of the evolutionary model, and *Κ* the substitution parameters. Moreover, P(D|G, Φ, *Κ*) is the observed sequence likelihood of an alignment D given the phylogenetic tree G, the collection of evolutionary rates (one for each branch) Φ, and the parameters governing the substitution model *Κ*. Here, f_T_(G | Θ) is the tree prior with hyperparameters Θ, f(Φ | Ω) the probability density of each branch’s evolutionary rate according to the latent model described below, and *π* (Ω, Θ, *Κ*) represents the prior over the remaining parameters.

In our model, Ω parametrizes the rate matrix underlying our latent-state CTMC and is comprised of two parameters, r and b, in addition to any parameters needed by the base evolutionary rate model – *µ* in the case of a strict clock. The rate matrix is defined as

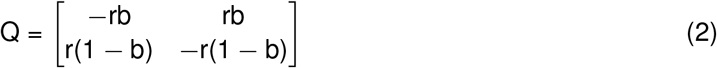

with the first state representing replication and the second latency.

Given that all nodes exist in the replicating state, the process on each branch is then independent of the others and so we write

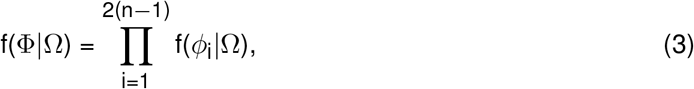

where n is the number of tips in our tree, 2(n − 1) is the number of branches and each branch has its own rate *ϕ*_i_. We define each 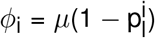 as the product of a base evolutionary rate *µ* and the proportion of the branch length spent in latency 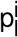. Here, we use a strict-clock model to assign the same base evolutionary rate to each branch, but more complicated models could be used. Thus, dropping the indexing for convenience, we write

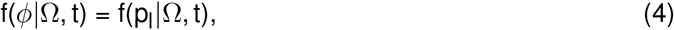

where t is the length of the branch in time. We derive f(p_l_|Ω, t) below.

We note that p_l_ = t_l_/t and so

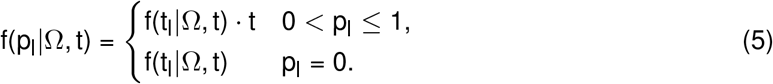

Let X(s) describe the state of a branch (i.e. replicating or latent) at time s (0 ≤ s ≤ t), and t_l_ describe the occupation time spent in the latent state. Sericola and colleagues (*60, 61*) provide numerically stable algorithms for the joint probability

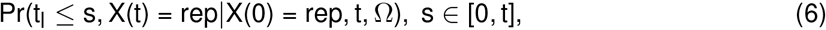

and also

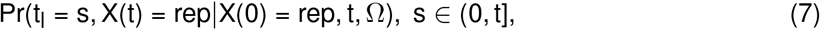

which are conditioned on the starting state being in the replicating state (X(0) = rep). These algorithms approximate an infinite sum to within a pre-defined error tolerance. Here we set the tolerance to 1 ×10^*−*10^.

Here, we are interested in conditioning on both the starting (X(0)) and ending (X(t)) state and have

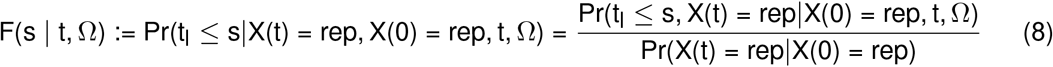

as the conditional cumulative distribution function (CDF) and

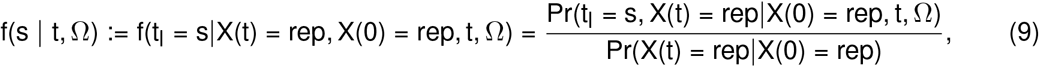

as the conditional density, which we will write with a t subscript below (F_t_(t_l_ |Ω, t) and f_t_(t_l_ |Ω, t)) to denote they are functions of the total branch length t. The probability of never entering the latent state t_l_ = 0 is given by F(0|Ω, t). The probability distribution function of the proportion of time spent latent f(p_l_|Ω, t) is then defined as

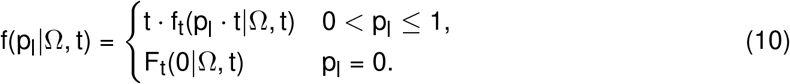

In our analyses we place a uniform (0, 1) prior on b and a modified conditional reference prior with rate equal to the height of the root on on r.

## Code availability

The latent-state model used above has been implemented in BEAST X (commit:edf73f) (*49*). All code needed to reproduce the analyses reported here has been deposited in Zenodo (https://doi.org/10.5281/zenodo.19225291) and can be found at https://github.com/ebov/latency-reservoir.

## Acknowledgments

JTM, MAS and AR are grateful for support from the Wellcome Trust (Collaborators Award 206298/Z/17/Z, ARTIC network). GB acknowledges support from the Research Foundation - Flanders (“Fonds voor Wetenschappelijk Onderzoek - Vlaanderen,” G098321N), from the European Union Horizon 2023 RIA project LEAPS (grant agreement no. 101094685), and from the DURABLE EU4Health project 02/2023-01/2027, which is co-funded by the European Union (call EU4H-2021-PJ4) under Grant Agreement No. 101102733. LMC was partly funded by FAPERJ - Fundação Carlos Chagas Filho de Amparo à Pesquisa do Estado do Rio de Janeiro, Processo SEI 260003/005679/2023 and SEI 260003/013252/2024. GD acknowledges the support of European Molecular Biology Organization (EMBO) installation grant IG-5305-2023. PM and AR acknowledge support from the Wellcome Trust (Discretionary Award 313694/Z/24/Z, ARTIC 2). IFO thanks the Wellcome Trust Hosts, Pathogens & Global Health program (Wellcome Trust, grant 218471/Z/19/Z) in partnership with Tackling Infectious Disease to Benefit Africa. MAS and AR acknowledge support from National Institutes of Health (NIH) grant R01 AI153044 and the European Research Council (grant agreement no. 725422 – ReservoirDOCS). MAS acknowledges further support from NIH grants R01 AI162611 and R01 AI192139.

## Supplemental information

### Supplemental Figures

**Figure S1:**
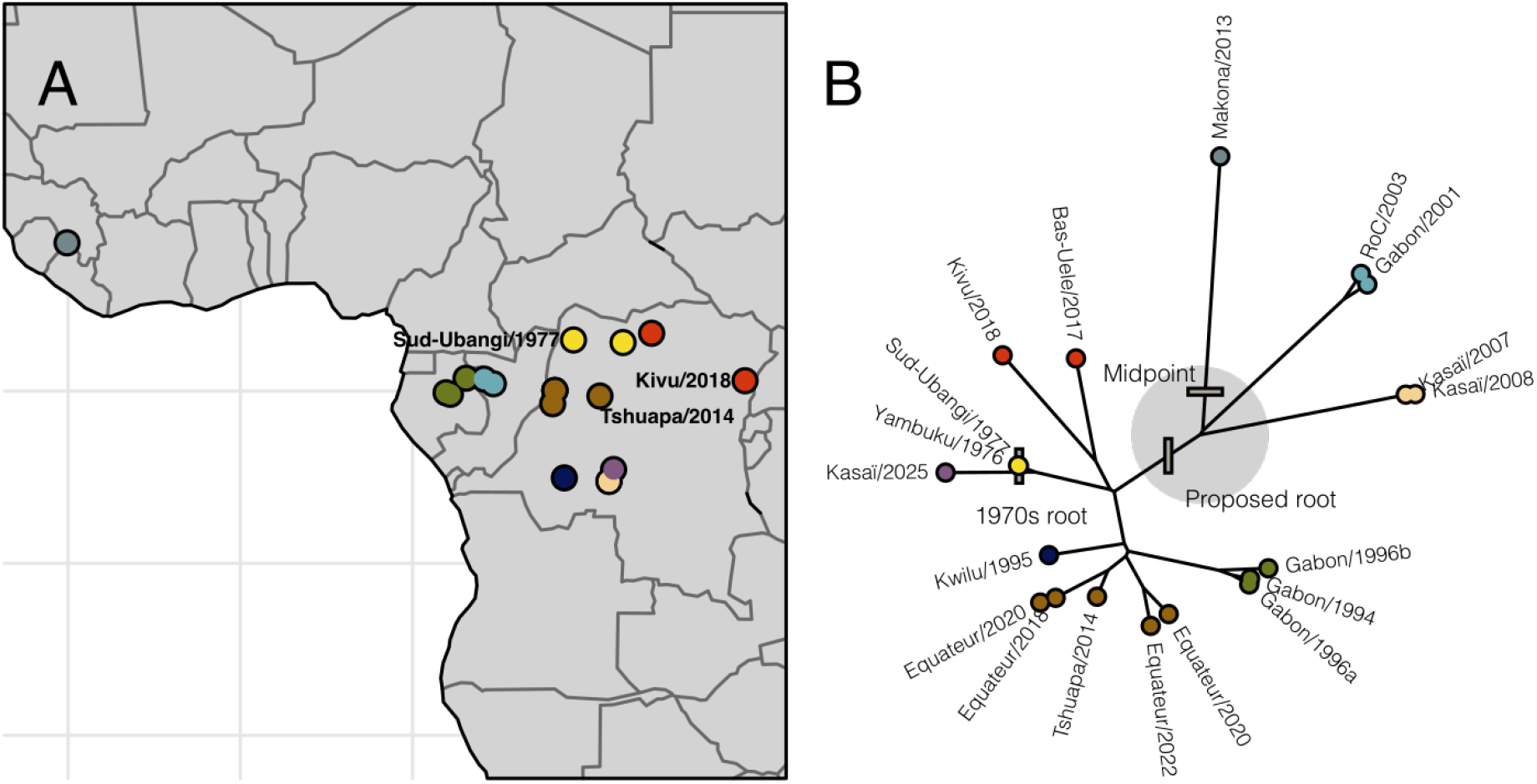
A map (A) and unrooted tree (B) representing the outbreaks included in this study. Common and proposed root placements are noted on the tree. The large grey sphere represents plausible root positions proposed by the latency rate model.

**Figure S2:**
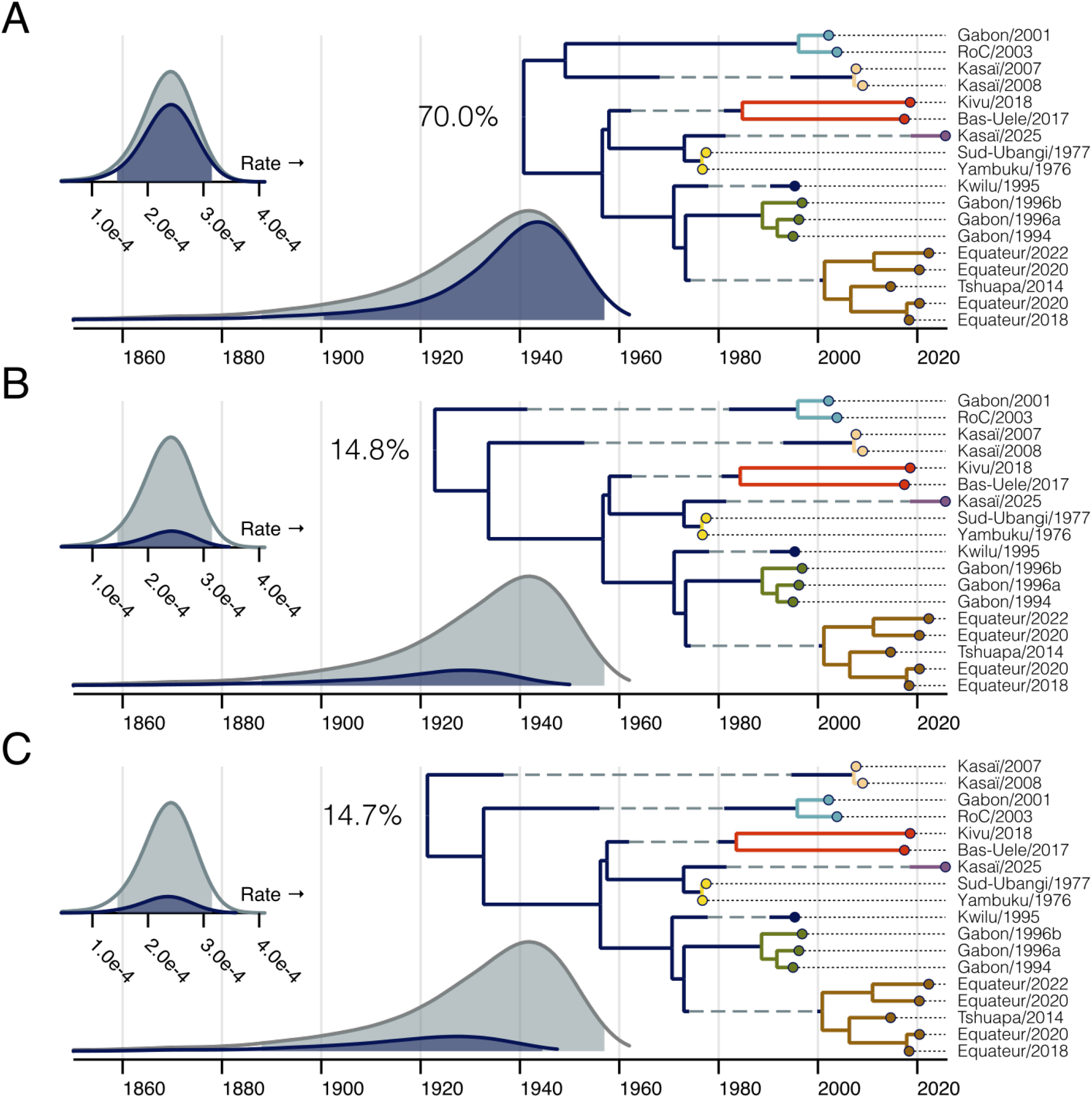
The evolutionary history of EBOV in Central Africa under the latent model (all explored roots). Same as Figure 2 but for all rootings with greater than 5% posterior support.

**Figure S3:**
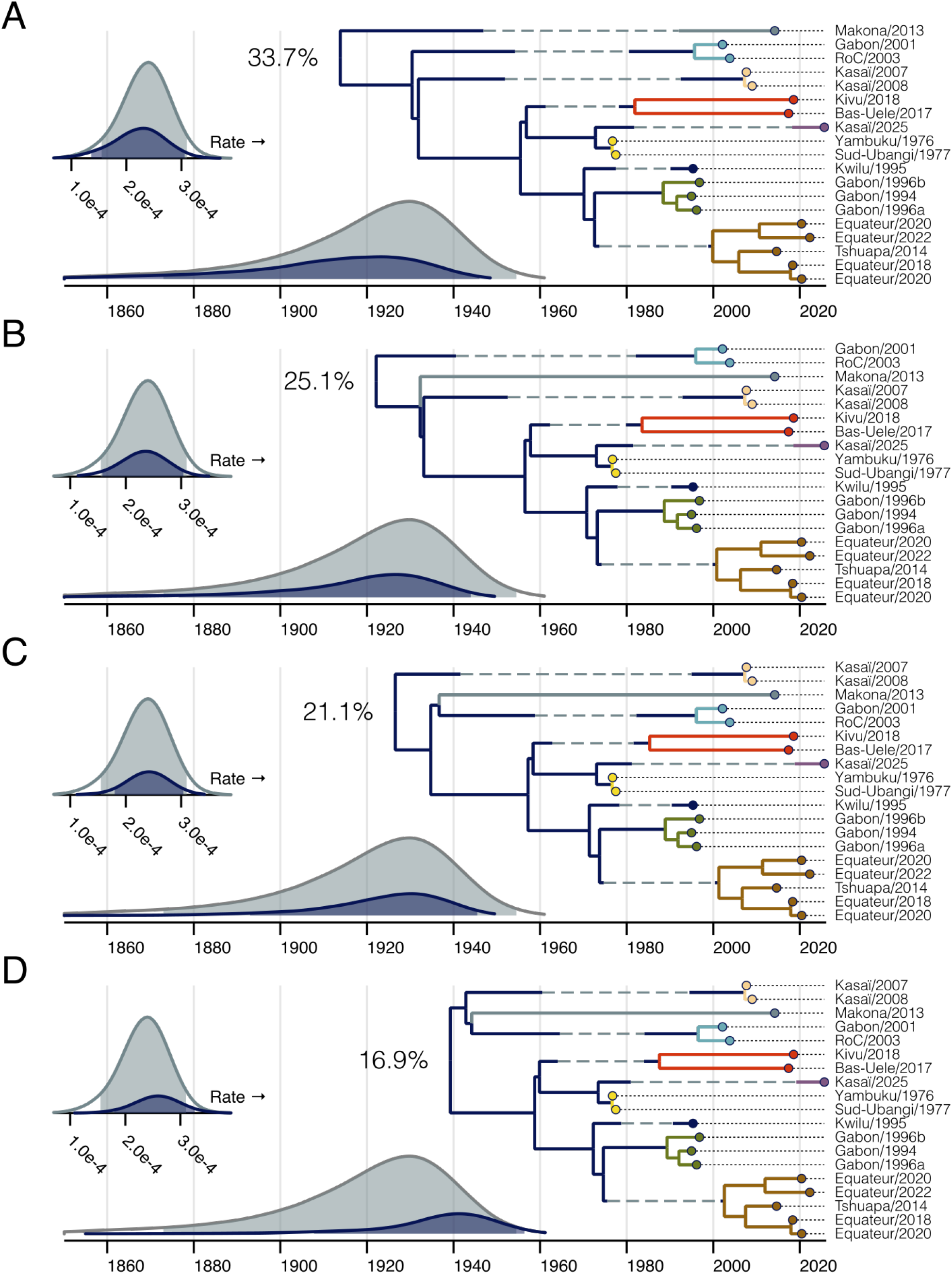
The evolutionary history of all EBOV outbreaks under the latent model partitioned by root position (all explored roots). Same as Figure 3 but for all rootings with greater than 5% posterior support.

**Figure S4:**
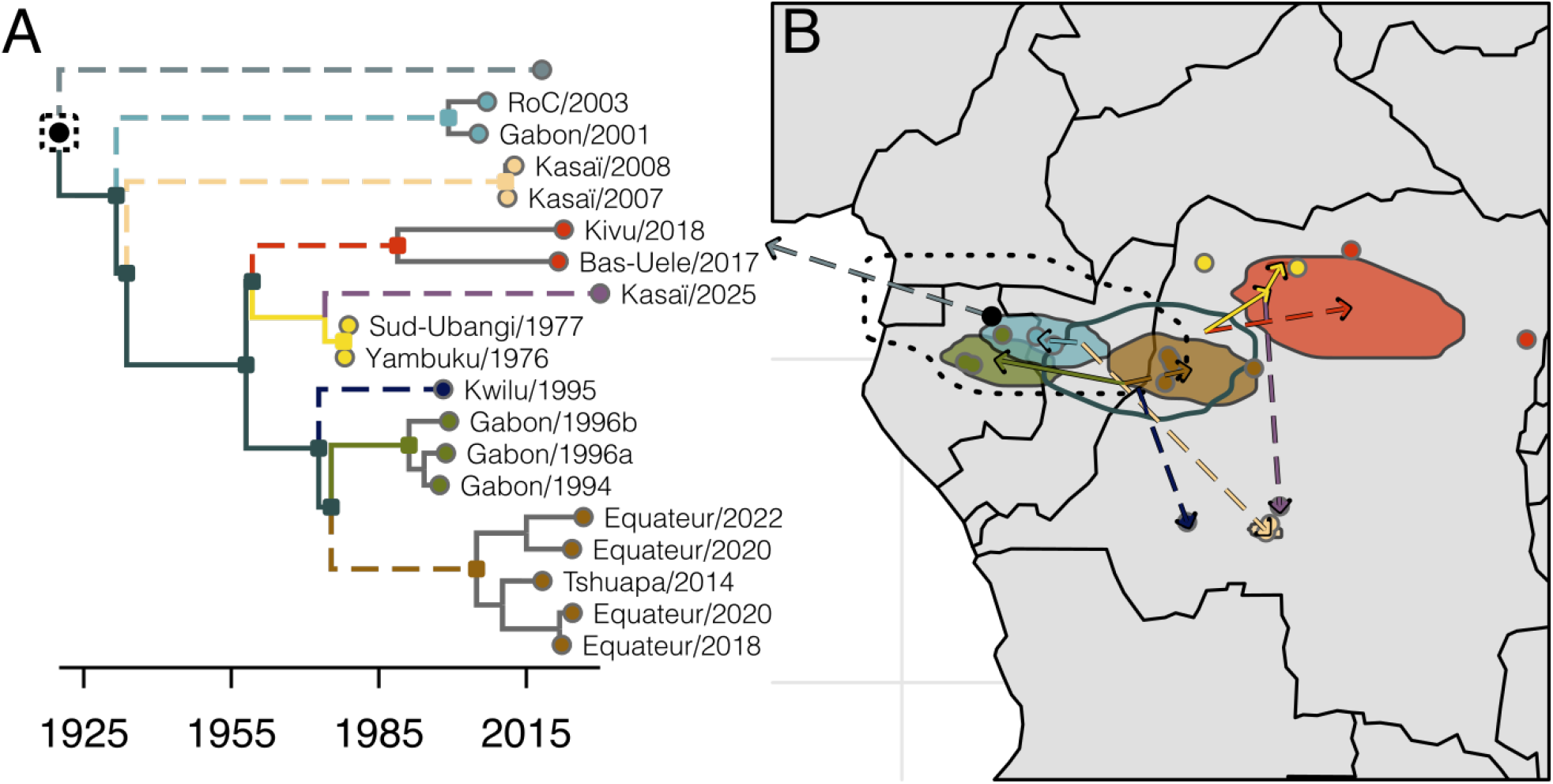
The geographic implication of latency in the EBOV reservoir including Makona/2013. Same as Figure 4 but the Makona/2013 sample has been included in the analysis.

**Table S1:** Recorded EBOV outbreaks used in this study. Outbreak - the name of the outbreak; Country-the country of the outbreak; Adm1 - Administrative Region 1 of the outbreak. Duration - the dates of the outbreak. * indicates two genomes from the Equateur/2020 outbreak are present in our data.

| Outbreak | Country | Adm1 | Dates |
| --- | --- | --- | --- |
| Yambuku/1976 | DRC | Mongala | Sep-Nov 1976 |
| Sud-Ubangi/1977 | DRC | Sud-Ubangi | Jun-77 |
| Gabon/1994 | GAB | Ogooue-Ivindo | Dec 1994-Feb 1995 |
| Kwilu/1995 | DRC | Kwilu | Jan-Jul 1995 |
| Gabon/1996a | GAB | Ogooue-Ivindo | Jan-Apr 1996 |
| Gabon/1996b | GAB | Ogooue-Ivindo | Jul-96 |
| Gabon/2001 | GAB | Ogooue-Ivindo | Oct 2001-Jul 2002 |
| RoC/2003 | COG | Cuvette Ouest | Oct-Dec 2003 |
| Kasai/2007 | DRC | Kasai Occidental | May-Nov 2007 |
| Kasai/2008 | DRC | Kasai Occidental | Dec 2008-Feb 2009 |
| Makona/2013 | GIN | Gueckedou | Dec 2013-Mar 2016 |
| Tshuapa/2014 | DRC | Tshuapa | Aug-Nov 2014 |
| Bas-Uele/2017 | DRC | Bas Uele | May-July 2017 |
| Equateur/2018 | DRC | Equateur | May-July 2018 |
| Kivu/2018 | DRC | Nord Kivu - Ituri - South Kivu | Aug 2018-June 2020 |
| Equateur/2020* | DRC | Equateur | June - Nov 2020 |
| Mbandaka/2022 | DRC | Equateur | April- July 2022 |
| Kasai/2025 | DRC | Kasai Occidental | August - October |

**Table S2:**
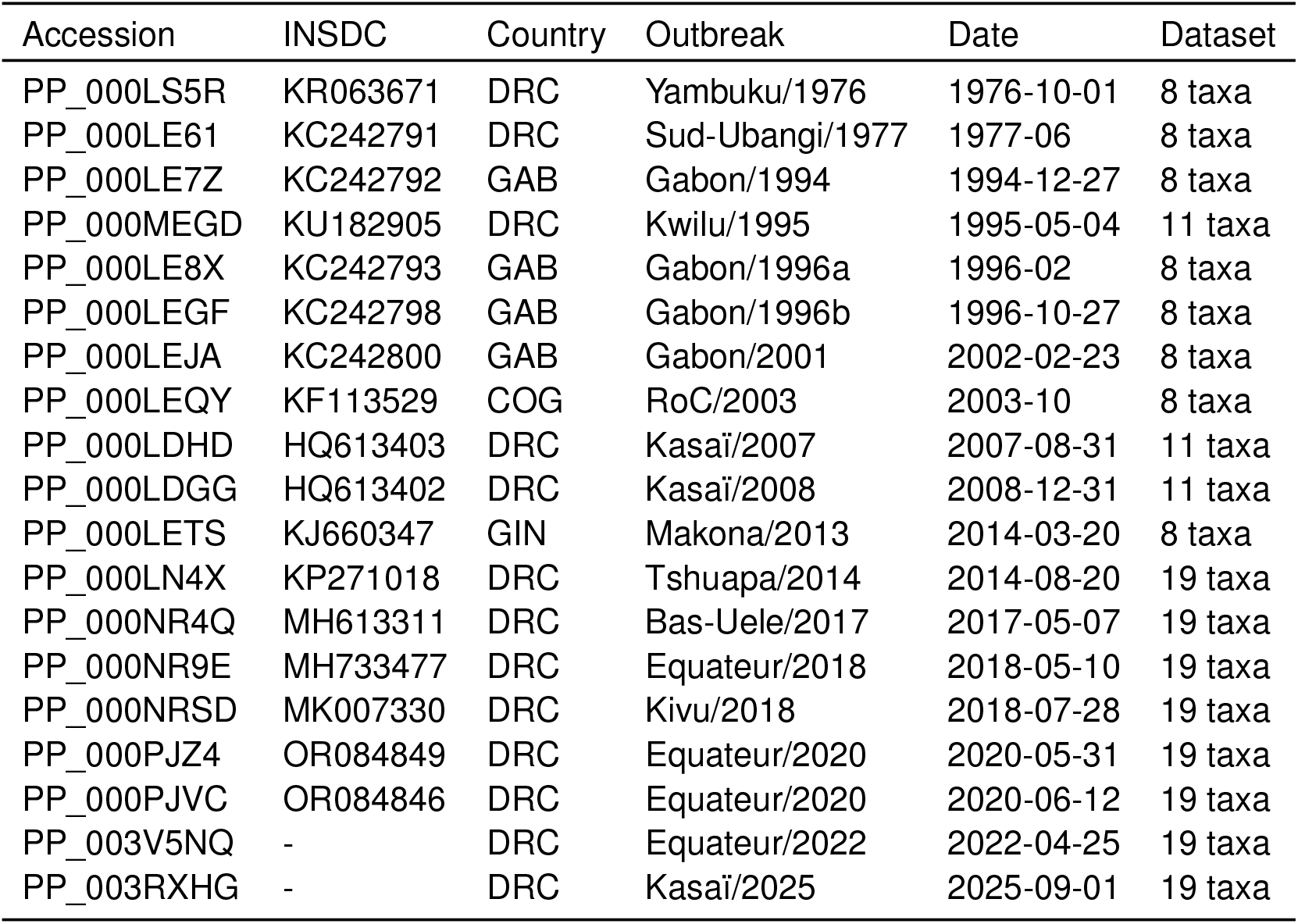
A list of the genomes used in this study with outbreak. Accession refers to Pathoplexus accession and can be found at (*41*). INSDC references the INSDC accesion for each sequence. Dataset denotes the smallest dataset in Table 1 which includes the taxa. Datasets build upon each other (e.g. the 11 taxa dataset adds 3 taxa to the 8 taxa dataset.)

## Supplemental text

### Further exploration of temporal signal

The temporal outliers observed in the EBOV phylogeny could have resulted from enhanced purifying selection instead of latency. To investigate this possibility, we repeated the analysis in Figure 1 with a partition made of only third position codon sites and intergenic regions (Figure S5). Despite elevated evolutionary rates, which are expected for a dataset enriched in synonymous sites, this analysis produced the same conclusions as that run on the full genome. Temporal outliers were maintained, as were our initial observations regarding within- and between-cluster rates.

### Prior sensitivity analysis

Preliminary analyses suggested that when applied to the EBOV dataset the latency model is positioned between two extremes, both of which likely have identifiability issues. If the rate of latent-replicating transition is too low relative to the scale of the tree, no latency is expected and the model reduces to a strict-clock model. In this case, there is no temporal signal in the full dataset if a strick-clock model is employed. The evolutionary rate is unidentifiable.

If the rate of transition is too high then the model approaches a free-rates model with an evolutionary rate on each branch. In some datasets where the parameters of the latency model may be known this could be tolerated. However, our analyses show that we have little power to predict *a priori* the amount of latency on a latent branch. Note the wide spread in the distribution of time spent latent in Figure 5B.

In this study we place a gamma prior on the rate of latent-replicating transitions. The gamma distribution was parameterized with a rate parameter equal to the root height. This prior was chosen to ensure the expected rate of latency scales with the phylogeny and strikes a balance between the extremes above without prohibiting either. As noted in the methods, this prior favors no latency while still allowing for the possibility of latency (95% prior probability of no latency).

**Figure S5:**
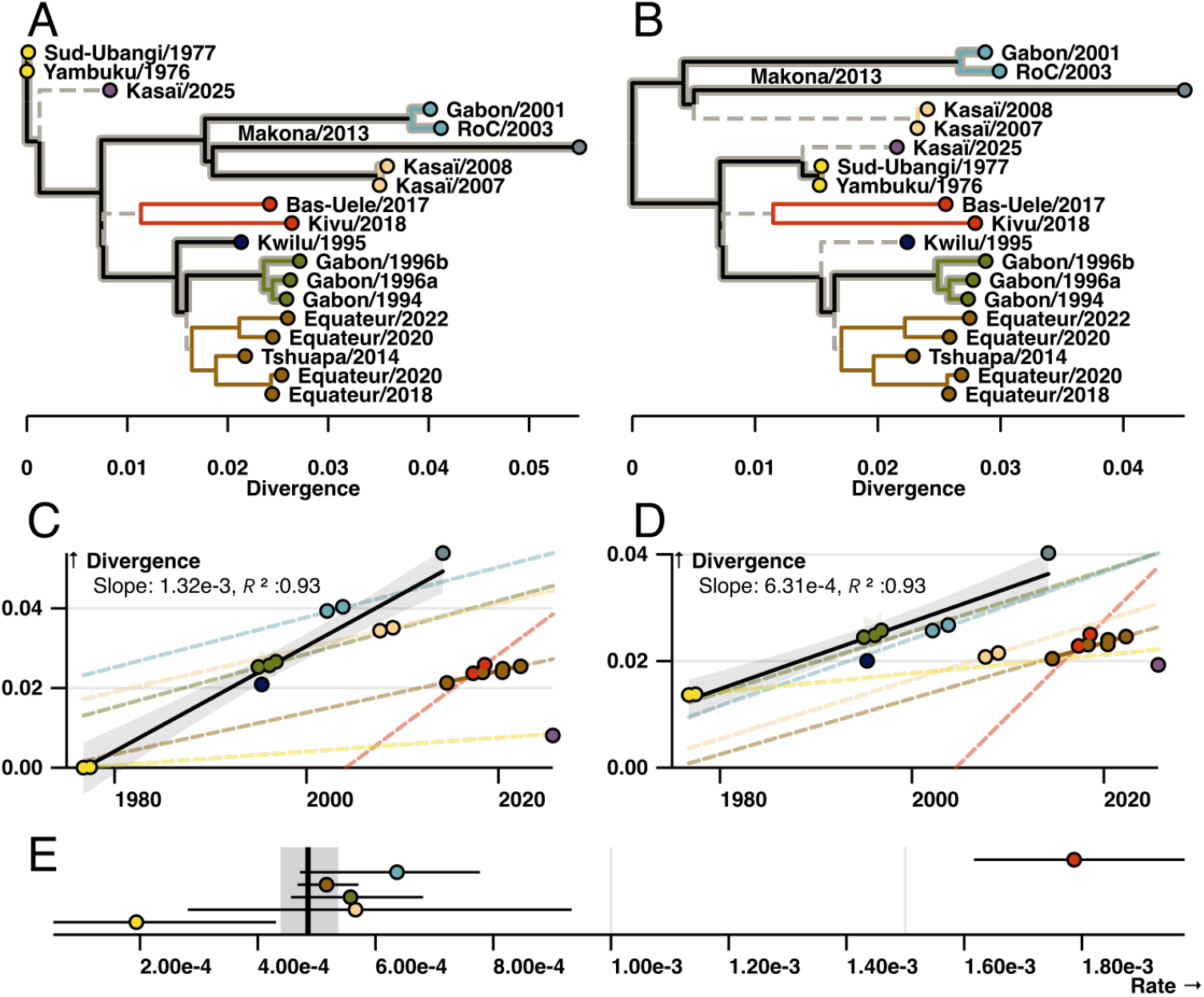
Rate heterogeneity in the EBOV phylogeny (third position coding sites and intergenic regions). Same as Figure 1 with the exception that the presented phylogenies were made with only sites in the third codon position and intergenic regions.

We ran two additional analyses to explore the impact of this prior on our findings. In the first, the rate of the gamma distribution was set to one half the root height, which would favor more latency (Figure S6). In the other, the rate of the gamma distribution was set to twice the root height, which favors less latency (Figure S7). Both analyses produced comparable results to that presented in the main text (Table S3 and Figures S7 & S6). In fact, increasing the rate of the gamma prior by a factor of two increased the posterior support for the proposed root to nearly 90% (Figures S7).

**Table S3:** Prior sensitivity analysis. Prior - the rate parameter of the gamma distribution placed on the latent rate. Root age-posterior mean and 95% HPD of the root age. Clock rate-posterior mean and 95% HPD of the evolutionary rate when replicating. Latent branches - the median and 95% HPD of the number of branches with at least one latent period.

| Prior | Root age | Clock rate ($\times 10^{-4}$) | Latent branches |
| --- | --- | --- | --- |
| Root height | 1929 [1887,1957] | 2.34 [1.41, 3.19] | 6 [5,9] |
| Root height $\div 2$ | 1924 [1878,1957] | 2.34 [1.42, 3.21] | 7 [5,11] |
| Root height $\times 2$ | 1934 [1903,1958] | 2.29 [1.36,3.08] | 6 [4,8] |

**Figure S6:**
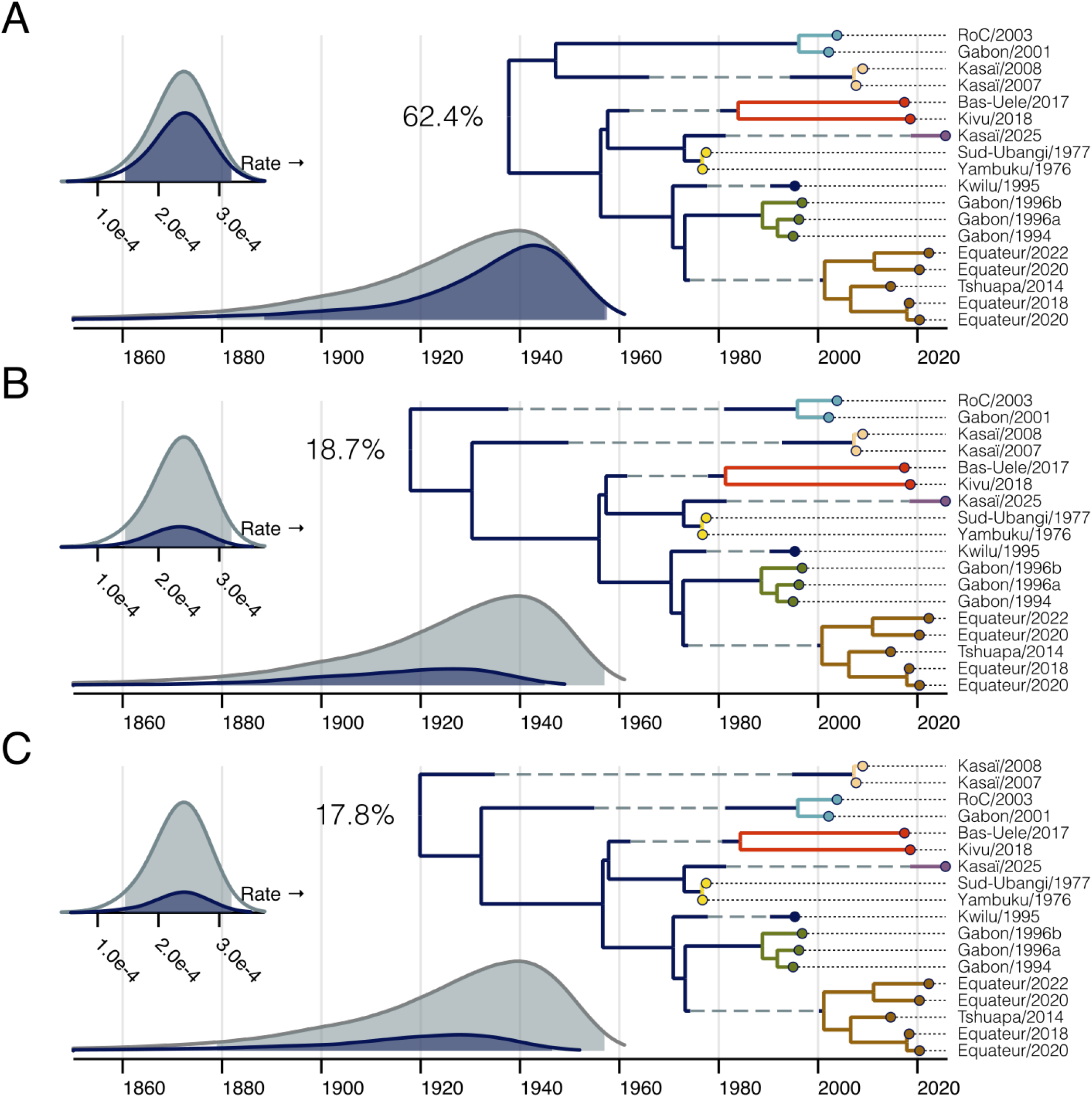
Evolutionary history of EBOV in Central Africa with a prior favoring more latency. Same as Figure S2; however the rate parameter of gamma prior placed on the rate of latent transitions is one-half the root height.

**Figure S7:**
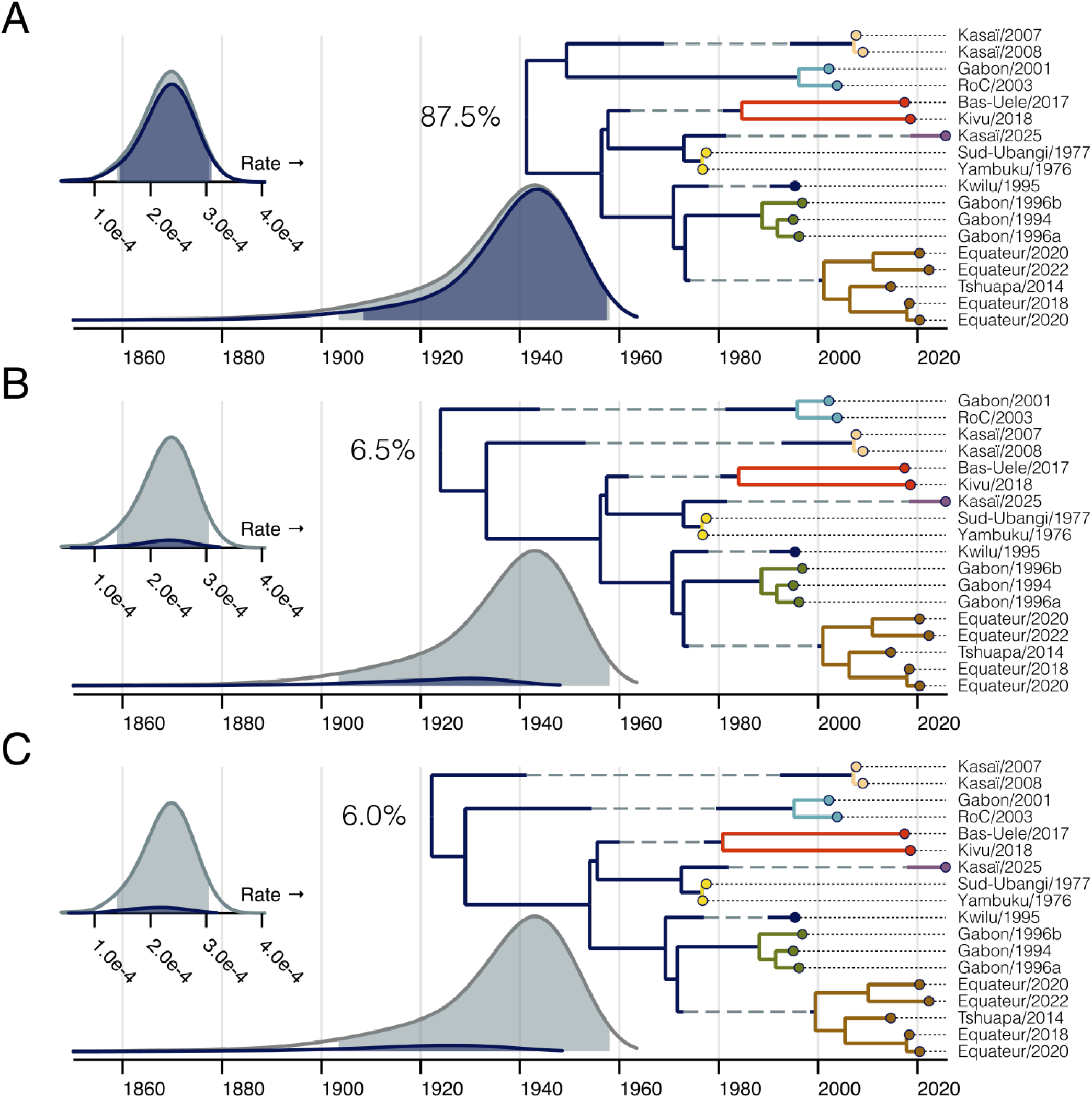
Evolutionary history of EBOV in Central Africa with a prior favoring less latency. Same as Figure S2; however the rate parameter of gamma prior place the rate of latent transitions is twice the root height.

